# Migratory birds are able to choose the appropriate migratory direction under dim yellow monochromatic light

**DOI:** 10.1101/2023.01.13.523666

**Authors:** Nadezhda Romanova, Gleb Utvenko, Anisia Prokshina, Fyodor Cellarius, Aleksandra Fedorischeva, Alexander Pakhomov

## Abstract

Previously it has been shown that migratory birds were oriented in the appropriate migratory direction under UV, blue and green monochromatic lights (short-wavelength) and were unable to use their magnetic compass in total darkness and under yellow and red light (long-wavelength). Currently, it is generally assumed that the magnetic compass of birds works correctly only under short-wavelength light. However, it also been suggested that the magnetic compass has two sensitivity peaks: in the short and long wavelengths, but with different intensities. In this project, we aimed to study the orientation of long-distance migrants, pied flycatchers (*Ficedula hypoleuca*), in different monochromatic lights during autumn migration. The birds were tested in the natural magnetic field (NMF) and 120° CCW shifted magnetic field (CMF) under green and yellow light (intensity 1 mW m^−2^). All tests were performed in a specially constructed wooden laboratory equipped with magnetic coils to manipulate the magnetic field. We showed that (1) pied flycatchers were completely disoriented under green light both in the NMF and CMF but (2) showed the migratory direction in NMF and the appropriate response to CMF under yellow light. Our data contradict results of previous experiments under monochromatic yellow light and might indicate the previously proposed hypothesis of two different mechanisms in avian magnetoreception (a high-sensitive short-wavelength mechanism and a low-sensitive mechanism in the long-wavelength spectrum) has a right to exist.

## Introduction

Currently, it is well known that migratory birds can detect the Earth’s magnetic field and use it as a cue source for orientation and navigation [1,2]. Even though the ability of birds to use information from the magnetic field for orientation was first described in the 1960s [3], the sensory, physiological and biophysical mechanisms of this compass system have still remained unexplored. One of the most popular theories, the radical pair model (RPM), proposed for the first time by Klaus Schulten [4] and lately upgraded by Thorsten Ritz and co-authors [5], assumes that animals have specialized magnetosensitive photoreceptors in the retina of the eye. According to this theory, the birds can perceive magnetic compass information through a process of light-dependent radical pairs in cryptochromes, light-sensitive proteins present in all cells of the animal’s body and the only known vertebrate photopigments that can form long-lived, spin-correlated radical pairs upon light absorption [5,6].

The avian magnetic compass has two main key features, which fit the RPM: the light dependence and the sensitivity to weak oscillating magnetic fields (OMFs) in the radiofrequency range. The initial radical pair theory predicts that an oscillating magnetic field in the lower megahertz range (1–100 MHz) can disrupt the magnetic compass due to the electron paramagnetic resonance effect [5,7]. Disruption of magnetic orientation in the presence of OMFs has been suggested as a diagnostic tool for the radical pair reaction mechanism underlying the magnetic compass in different animal taxa [8–10], but see [11]. This effect of OMFs was experimentally observed in various studies performed by independent scientific groups in migratory [12–16] and non-migratory birds [17,18]. However, it should be noted that this model cannot fully explain the results of some behavioural studies: 1) the sensitivity thresholds of the magnetic compass to OMFs in European robins and garden warblers are two orders of magnitude lower than what the mainstream theory predicts [19–22]; 2) there is no consensus among researchers whether the magnetic compass orientation of birds can be disrupted by both narrow-band (at the Larmor frequency) and broadband electromagnetic fields or only by broadband electromagnetic noise [14,15]; 3) a recent study showed the insensitivity of the avian magnetic compass to OMFs applied locally to the eyes [22].

Magnetic compass orientation in birds (migratory and non-migratory) and several species of other animal taxa (butterflies, newts, frogs, beetles, and fruit flies) has been shown to be light-dependent [17,23–30]. Results of various behavioural studies indicate that both the spectrum wavelength and light intensity play a crucial role in the ability of migratory birds to orient using information from the Earth’s magnetic field. Garden warblers *Sylvia borin*, Australian silvereyes *Zosterops lateralis* and European robins *Erithacus rubecula* are able to choose the appropriate migratory direction in orientation cages under full-spectrum (‘white’) light and different types of monochromatic light from UV to green light (short-wavelength light, 373-565 nm) and show disorientation or a shifted response under long-wavelength light (from yellow to red, 590-645 nm) or total darkness [23,31,40–46,32–39] (for more details about response of tested birds to different parameters of monochromatic light in behavioural studies, see Table S1). The intensity (irradiance mW/m^2^ and photon quantity) is the next crucial characteristic of light that affects the magnetic orientation of birds. It has been shown that high light intensity interferes with the ability to use properly the magnetic compass for orientation: birds were able to choose the appropriate migratory direction when tested under low-intensity UV (0.3 mW m^−2^, 0.8×10^15^ quanta s^−1^ m^−2^; [46]), turquoise (2.1 - 2.2 mW m^−2^, 7 - 8×10^15^ quanta s^−1^ m^−2^; [38,43,47]), blue (2.4 - 2.8 mW m^−2^, 8 - 8.7×10^15^ quanta s^−1^ m^−2^ [33,38,40,46,47] and green (0.2 - 3 mW m^−2^, 6 - 8.7×10^15^ quanta s^−1^ m^−2^ [22,23,32–36,42,47]) monochromatic light. If the light intensity inside orientation cages was increased (2-5 fold or more, depending on the spectrum of light), birds were disoriented or showed an axial response (for details, see Table S1).

However, the relation between the avian magnetic compass and the parameters of light is more complex than described above. In the absence of light (‘total darkness’), birds are active and able to orient but in a direction that does not correspond to the population-specific migratory direction and is independent of the migratory seasons (the same in autumn and spring; [44,45]). According to the authors of this study, birds do not have the opportunity to use the light-dependent inclination magnetic compass under such light conditions and show a so-called ‘fixed direction’ response [44]. It is based on a magnetite-based magnetic polarity sense: 1) birds altered their headings accordingly when tested in the magnetic field with the horizontal component reversed and do not respond to the inversion of the vertical component; 2) OMFs had no disruptive effect on such behaviour of birds in orientation cages; 3) “fixed direction” responses are disrupted when the birds’ beaks are treated with a local anaesthetic Xylocaine (lidocaine) which can deactivate putative magnetite-based magnetoreceptor in the upper beak ([2,48] but see [49]). The 1 hour pre-exposure to a light condition other than full-spectrum light (total darkness and green light) affects the orientation behaviour of the tested birds and leads to disorientation or axial response when the birds are tested in green light, in contrast to blue or turquoise light ([47], but see [42]).

Adding dim yellow (590 nm, 2.0 mW m^−2^) to blue (424 nm, 2.8 mW m^−2^) or green (565 nm, 2.1 mW m^−2^) light results in an unusual behaviour of birds in orientation tests: they show direction which does not correspond to the normal seasonal migratory direction and differs between seasons [41], similar to orientation in total darkness. The authors of this study suggested that changes in orientation of birds under blue-yellow and green-yellow light were caused by interaction between two different receptors (short-wavelength and long-wavelength). Previously, a similar idea was proposed in another research carried out during autumn migration: European robins were oriented west-northwest under low-intensity red light (1mW m^−2^, 3.2×10^15^ quanta s^−1^ m^−2^; 5mW m^−2^, 16×10^15^ quanta s^−1^ m^−2^), while at high intensity (10 mW m^−2^, 32×10^15^ quanta s^−1^ m^−2^) they were disoriented [39]. It allows us to assume that birds might have a light-dependent magnetoreception system based on two spectral mechanisms: a highly sensitive short-wavelength mechanism in the blue-green spectrum and a low-sensitive long-wavelength mechanism in the red spectrum. The existence of a similar system was observed in red-spotted newts *Notophthalmus viridescens*: they exhibited shoreward (normal) magnetic orientation under full spectrum light/ 450 nm light and 90° counterclockwise shifted orientation under light with wavelengths > 500 nm [25,50]. However, independent replication of this study in European robins and Australian silvereyes did not confirm these findings [45]: both species were oriented in a similar direction under dim red light (1mW m^−2^) as robins in the previous study, but this direction did not differ significantly between migratory seasons (spring and autumn) and light conditions (red light and total darkness). All of these indicate that it might be a ‘fixed direction response discovered previously by the same scientific group [40,44].

Despite all the exceptions mentioned above, currently, it is generally accepted that birds can orient using information from the geomagnetic field only under short-wavelength light (from UV to green) and lose this ability in long-wavelength light (yellow and red). On the other hand, it is difficult to fully ignore the fact that orientation in long-wavelength light might be possible according to the results of behavioural studies in birds and other species that have not yet been fully explained [25,26,39,50,51]. In the present study, we aimed to investigate magnetic compass orientation in a new model species, long-distance migrant pied flycatcher *Ficedula hypoleuca*, under dim green and yellow light, which are on the “threshold” between the short-wavelength and long-wavelength light.

## Materials and methods

### Study site, model species and bird keeping

The capture of birds and experiments were carried out in the vicinity of the Biological Station Rybachy on the Curonian Spit (Kaliningrad Region, Russia; 55°09′N, 20°52′E) in August-September 2021-2022. As a model species, we used first-year pied flycatchers (male and female), common migrants, and a new object to study light-dependent magnetoreception in migratory birds. After capture by mist-nets, birds were placed in a laboratory aviary without access to any astronomical cues (stars and the sun) under an artificial photoperiod that corresponded to the natural one and was controlled by an IoT dimmer Shelly Pro 1 (Allterco Robotics, Bulgaria). Each bird was kept in a separate compartment of a cage (40×40×40 cm) and fed mealworms, mixed diet (eggs, carrot, breadcrumbs), and Padovan complete feed for insect-eating birds + a vitamin supplement in pure water *ad libitum*. Fat score and weight of each bird were estimated every day for 3-4 days after trapping. If these parameters (fat and weight) constantly decreased for two days (which indicates a too high level of stress), we released these birds into the wild. The indoor aviary was equipped with IP cameras with infrared LEDs (840 nm) so that we could monitor the activity and behaviour of the birds in their cages in real time at night. For experiments, we selected only birds which exhibited migratory restlessness on a given night.

All animal procedures (the capture of the birds and simple, non-invasive, behavioural experiments) were approved by the appropriate authorities: Permit 24/2018-06 by Kaliningrad Regional Agency for Protection, Reproduction and Use of Animal World and Forests; and Permit 2020-12 by Ethics Committee for Animal Research of the Scientific Council (Zoological Institute, Russian Academy of Sciences). All birds were released at the end of all tests well before the migration of their conspecifics finished.

### Experimental setup and test conditions

Before each test, the birds were kept in darkness for at least an hour to avoid the cumulative effect in cryptochromes. According to the previous study, birds can use products of the first step in the redox cycle of FAD that accumulate before the light is turned off. If you then test birds under artificial light conditions (darkness, monochromatic lights, etc.), it can lead to a false directional result [52].

All birds were randomly tested under different light and magnetic experimental conditions: 1) green light and the natural magnetic field (NMF); 2) green light and a changed magnetic field (CMF) with magnetic north deflected 120° counterclockwise (120° CCW); 3) yellow light and the NMF; 4) yellow light and 120° CCW CMF. Experimental magnetic fields were produced by a double-wrapped, three-dimensional Merritt four-coil system (‘magnetic coils’ hereafter). This Merritt coil system generates a field with >99% homogeneity within a space of ca. 110 × 110 × 110 cm. Each of the three axes of coils was driven by a separate constant current power supply BOP 50-4M (Kepco Inc., USA) placed along with the coils’ control box in a shielded and grounded box, located about 2 meters from the Merritt coil system. The box reduced the sound of working power supplies so it was by far below the level of natural environmental noise near the experimental setup and could not be an audible position cue for tested birds. We did not find any effect of working power supplies on bird orientation according to the results of our previous studies in which the same power supply box was used [53–55]. The parameters of the magnetic field were checked in the centre of the experimental table and opposite corners using a FMV400 portable magnetometer (Meda Inc., USA) before and after each test night. The average magnetic measurements over all test days under the NMF condition: intensity 50463 ± 104 nT (mean ± sd) and inclination 70.11° ± 0.55; under the CMF condition: intensity 50468 ± 105 nT, inclination 70.21° ± 0.42° and declination (horizontal direction) -119.97° ± 0.11°.

Behavioural experiments under artificial light conditions took place in a non-magnetic laboratory chamber specially constructed around the magnetic coils (Figure S1). We used professional LED strips (Arlight, Russia) as a light source of monochromatic lights (see parameters and spectrum of LEDs in Figure S2 and Table 1). LED strips were mounted on an aluminium frame (under the chamber ceiling, Figure S1B) and fed by a Rigol DP711 programmable DC power supply (Rigol Technologies, Inc., Beaverton, USA) which was placed in the power supply box (mentioned above). The LI-1500 lightmeter (LI-COR Inc., USA) with a LI-190R terrestrial quantum sensor was used to measure the level of light intensity (in quanta). Measurements in irradiance (mW m^−2^) under monochromatic lights were taken by the P-971-1 optometer (Gigahertz Optik, Germany) and the probe ‘Visible’ RW-3703-2, a silicon photo element for the wavelength range 400-800 nm (with specific calibrations for the wavelengths of the LEDs; [52]), in full spectrum natural light (outside) and ‘darkness’ (inside) were taken by the LI-1500 lightmeter with the LI-200R pyranometer sensor. We measured the spectrum of LEDs using the MK350N PREMIUM handheld spectrometer (UPRtek Inc., Taiwan). All measurements of light intensity and spectrum have been taken inside the orientation cages under Plexiglass lids, at the level of the birds’ head.

**Table 1.** Light conditions in orientation cages during experiments. Light intensities are given in radiometric irradiance (mW m^−2^) and photon quantities (quanta s^−1^ m^−2^), m – monochromatic light condition.

| Monochromatic light | The peak wavelength, nm | Half bandwidth, nm | Range of half bandwidth, nm | Irradiance, $\text{mW m}^{-2}$ | Photon quantities, $\times 10^{15} \text{ quanta s}^{-1} \text{m}^{-2}$ |
| --- | --- | --- | --- | --- | --- |
| Green | 530 | 29 | 517 - 546 | 1 | 2.9 |
| Yellow | 589 | 16 | 580 - 596 | 1 | 4.2 |

Parallel / before the experiments under monochromatic lights in the laboratory chamber (see dates of experiments under each experimental condition in Table S1), the birds were tested under natural light conditions (full-spectrum light) with (autumn season 2021) and without access to stars (autumn seasons 2021-2022) to make sure that our birds were in migratory condition and could show species-specific migratory direction in orientation cages. All experiments under natural light condition were performed outdoors on wooden tables placed in the clearing of reeds on the coast of the Courish Bay near the laboratory chamber. This experimental place is in a rural location with low levels of radio-frequency noise according to our previous [14] and recent measurements of RF performed in 2022 (unpublished data) using a calibrated active loop antenna FMZB 1512 (Schwarzbeck Mess-Elektronik, Germany) and a spectrum analyzer Rigol DSA815-TG (Rigol Technologies, USA). In 2022 after the birds were tested under the monochromatic light condition in the NMF and the CMF, they were tested again under the NMF condition in so-called “darkness” (when all light sources inside the laboratory chamber were turned off).

### Orientation tests and data analysis

All orientation experiments (in full-spectrum and monochromatic lights) started at least 1 hour after local sunset (usually at the beginning of astronomical twilight, when the all tested birds started to show nocturnal restlessness in cages) and lasted 25-30 min for each experimental condition. We used Emlen funnels (the diameter of the upper part - 34 cm, the lower part - 10 cm, the height - 14 cm, the angle of the wall-45°, material – aluminium; [56]) to analyze the direction of birds’ activity under different lights. In all experiments (except tests in full-spectrum light + stars) on top of Emlen funnels, we put lids made of opaque glass (Plexiglass XT, 3 mm, 70% transmission) which completely obscured stars and any other patterns (landscapes, magnetic coils, etc) and diffused light from LEDs. Thus, the only orientation cue available to experimental birds during the test was the Earth’s magnetic field.

The directionality of the birds’ activity was recorded as scratches left by their claws as they hopped in the funnels on a print film covered with a dried mixture of whitewash and glue. The data (each bird’s mean direction from the distribution of scratches) were analysed by 3 independent researchers (2021: N.R., A.F., Dmitry Sannikov; 2022: A.Pr, Dmitry Sannikov, Julia Bojarinova), who did not know under which magnetic conditions (CMF or NMF) a particular bird was tested. In most cases, the mean direction could be very precisely identified using the simple visual estimation method [57]. If a pattern of scratches was not clear, the scratches were counted in each of 36 10 ° sectors, and we used a custom-written script to assess the directionality based on the number of scratches. The mean of the directions determined directions was recorded as the orientation result. If at least two observers considered the scratches to be randomly distributed or if the two mean directions deviated by more than 30º, the bird was considered to be not oriented in the given test. If the number of scratches was fewer than 40 scratches, the bird was considered to be inactive in a given test.

For the final analysis, we included the results of the birds tested at least two times with at least one result significantly oriented according to the Rayleigh test [58] at the 5% significance level. We performed additional distribution analysis (maximum likelihood analysis) for our samples using the CircMLE R package, as it was described in [59], to be fully sure that our birds showed unimodal orientation under different experimental conditions. The nonparametric Mardia-Watson-Wheeler (MWW test) test was used to test the significance of the difference between the orientation of pied flycatchers under different conditions. The data used for the final analysis are accessible in Table S2 in the Supplementary Materials. Circular statistics were performed in Oriana 4.02 (Kovach Computing Services, UK). Additionally, we used the bootstrap technique [60] to identify whether significantly oriented groups showed significantly more directed behaviour than non-statistically significantly oriented groups. According to this method, a random sample of orientation directions (n angles) was drawn with replacement from the sample of orientation directions present in the significantly oriented group. Based on these n orientation angles, the corresponding p-value was calculated, and this procedure was repeated 100 000 times [13,61]. After that, the resulting 100,000 R-values are ranked in ascending order: the r-values at rank 2500 and 97500, at rank 500 and 99500 define the 95% and 99% limits for the actually observed r-value of the significantly oriented group, respectively. If the actual observed r-value of the disoriented group is outside these confidence intervals, the oriented group is significantly more directed than the disoriented group with a significance of p < 0.05 and p < 0.01, respectively. Circular statistics were performed Oriana 4.02 (Kovach Computing Services, UK). Bootstrap and maximum likelihood analysis were performed in R 4.1.1 [62].

## Results

Under full spectrum (natural) light pied flycatchers showed the seasonal population-specific migratory direction in southwest without the opportunity to obtain directional information from astronomical cues (under plexiglass lids, 2021-2022: α = 212°, n = 28, r = 0.491, p < 0.001, 95% CI=183°-241°, Fig. 1A) and with access to stars (2021: α = 212°, n = 18, r = 0.643, p < 0.001, 95% CI=187°-237°, Fig. 1B). The mean direction of birds obtained in the control tests under natural light was similar to the mean autumn migratory direction of the same species on the Curonian Spit according to ringing data [63] and previous laboratory and field studies [53,64].

**Figure 1.**
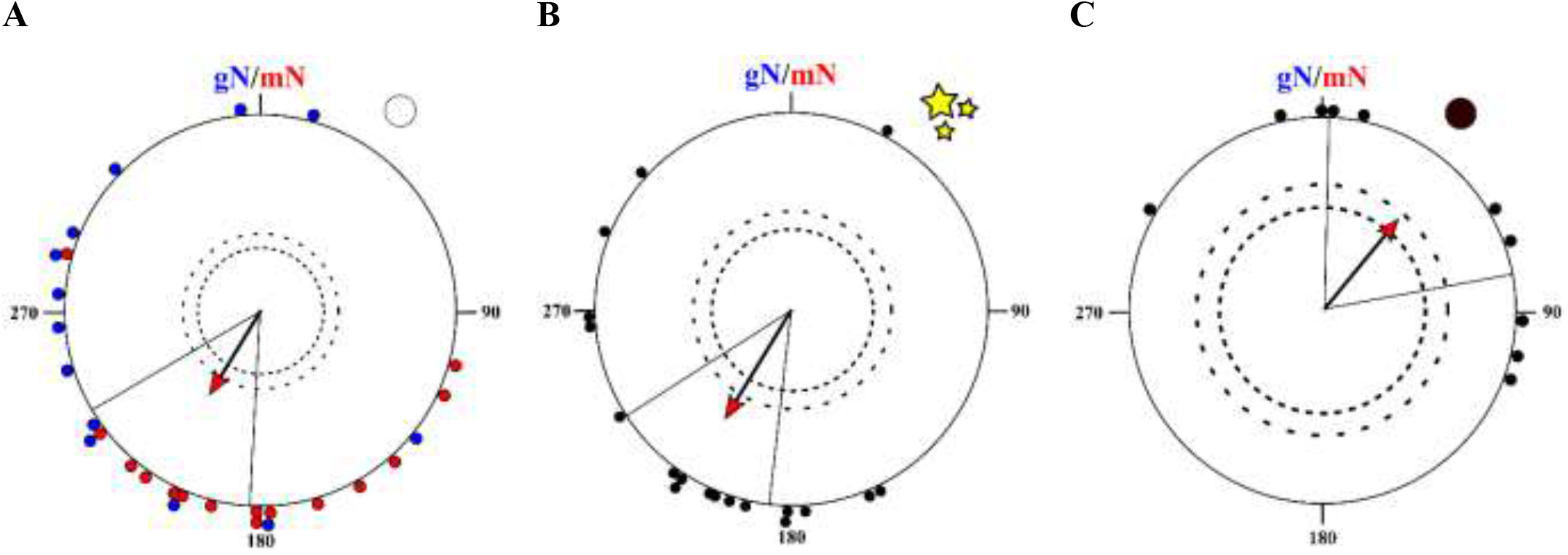
Orientation of juvenile pied flycatchers under different full-spectrum light conditions and total darkness: A) outdoor experiments without access to stars (under Plexiglass lids), 2021-2022; B) outdoor experiments with access to stars, 2021; C) indoor experiments in “darkness”, 2022. Each dot at the circle periphery indicates the mean orientation of one individual bird; the colour dots in A indicate orientation of birds in different years: 2021 (red) and 2022 (blue). The inner and outer dashed circles represent the 1 and 5 % significance levels of the Rayleigh test, respectively. Geographic (gN) and magnetic (mN) Norths correspond to 0° (local declination is 6°), and radial lines indicate the 95 % confidence intervals [CI] for the group mean orientation direction.

Experiments in monochromatic green light showed that the birds could not determine the migratory direction either in the natural (α = 115°, n = 10, r = 0.199, p = 0.68; Figure 2A) or in 120° CCW rotated magnetic fields (α = 238°, n = 12, r = 0.168, p = 0.72; Figure 2B). Results of the bootstrap analysis indicate that direction chosen by birds under green light conditions were significantly more scattered than in NMF under yellow light in 2021 (NMF_yellow_2021 - NMF_green_2021: p < 0.01; 95% CI for r-value is 0.48 < r <0.86; 99% CI for r-value is 0.44 < r <0.94, Fig S3A) and in CMF under yellow light (CMF_yellow_2022 - CMF_green_2022: p < 0.01; 95% CI for r-value is 0.42 < r <0.93; 99% CI for r-value is 0.38 < r <0.94, Fig S3B).

**Figure 2.**
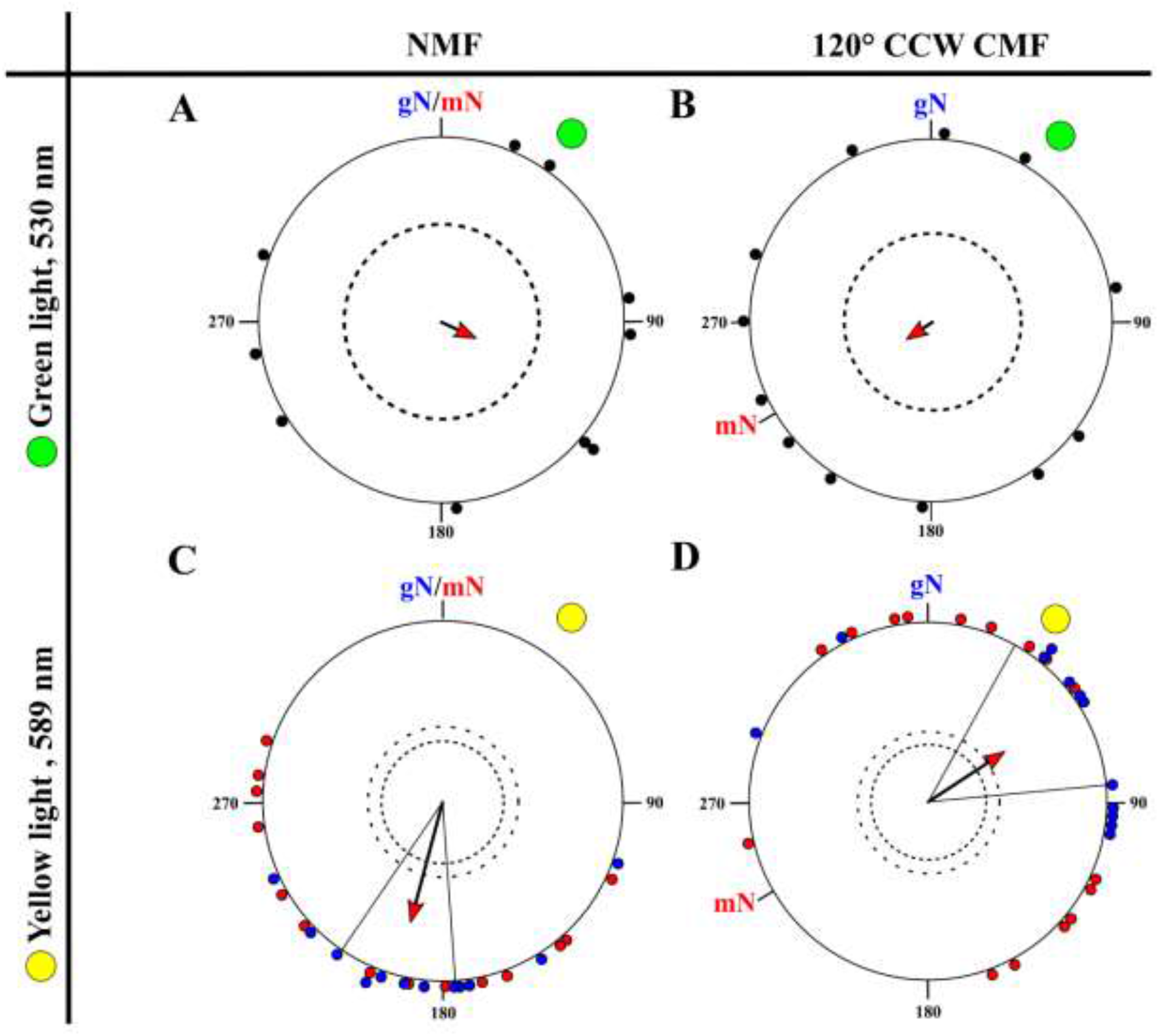
Orientation of juvenile pied flycatchers under different monochromatic light conditions: A) green light, the natural magnetic field (NMF), 2021; B) green light, 120° counterclockwise rotated magnetic field (120° CCW CMF), 2021; C) yellow light, the natural magnetic field (NMF), 2021-2022; D) yellow light, 120° counterclockwise rotated magnetic field (120° CCW CMF), 2021-2022. gN – geographic North, mN – magnetic North. For a description of the circular diagrams, see the legend of Figure 1.

Surprisingly, under monochromatic yellow light, the birds showed the seasonally expected migratory direction under NMF condition (α = 195°, n = 26, r = 0.7, p < 0.001, 95% CI = 177° - 214°, Figure 2C). The orientation under this experimental condition was not significantly different from the orientation of the same birds in similar magnetic condition under natural full-spectrum light (the MWW test: W = 1.94, p = 0.38 and both 95 and 99% confidence intervals of these distributions overlap: 99% CI in natural light: 174° – 250°, 99% CI in yellow light: 171° – 220°). When we turned the horizontal component of the magnetic field by 120° counterclockwise, pied flycatchers apparently responded to such magnetic field manipulation (α = 57°, n = 28, r = 0.5, p < 0.001, 95% CI = 29 - 85, Figure 2D). The orientation of pied flycatchers under the CMF condition under yellow light was significantly different from the orientation of the same birds in NMF both under yellow and full spectrum light (the MWW test: W = 31.2, p << 0.001 and W = 25.4, p << 0.001, respectively), their 95 and 99% do not overlap (99% CI in CMF yellow light: 20° – 93°) and both confidence intervals (95 and 99%) under the CMF yellow light condition include the expected 120° CCW rotation relative to mean direction in the NMF (α = 75°). Under “darknes” condition in 2022, the birds showed northestern direction (α = 39º, n = 10, r = 0.61, p = 0.02, Figure 1C) which does not correspond with autumn migratory direction typical for this species. According to the results of the maximum likelihood analysis, birds showed unimodal orientation under the majority of experimental conditions (Table S3), with the exception in darkness, where a bimodal alternative is suggested, probably due to a small sample size.

## Discussion

Experimental data obtained in this project clearly show that long-distance songbird migrants, pied flycatchers, are not able to use their magnetic compass under low-intensity green monochromatic light. These results contradict the generally accepted view that light-dependent magnetoreception in migratory birds takes place only under full-spectrum or monochromatic light from UV to green [65,66]. In our experiments, we used green LEDs with a peak wavelength of 530 nm which was in the middle between turquoise (502-510 nm) and green (560-571 nm) light used in previous studies [23,38,39,43]. However, as mentioned in Methods, the birds were transported (in opaque textile bags) from the windowless aviary to the temporary wooden house (on the experimental site) at least 1 hour after the lights in the aviary were turned off. Under this condition, they did not have the opportunity to use the products of the first stage in the redox cycle of FAD that accumulated under full-spectrum or UV/blue light [47,52]. Flavin adenine dinucleotide (FAD) is the light-absorbing cofactor of cryptochromes (its ‘light antenna’) and exists in three different redox states with different absorption spectra (FAD_ox_, FADH^·^ and FADH^−^). During the first stage of the redox cycle, the fully oxidised state, FAD_ox_, reduces to the neutral semiquinone radical FADH^·^ after absorption of full-spectrum or monochromatic light with wavelengths from UV to blue (peaks of absorption at 360 and 470 nm) [66,67]. FADH^·^, which forms a magnetic sensitive radical pair with a tryptophan radical [68], can be further light-independently reoxidised to FAD_ox_ or photoreduced by UV - green light (peaks of absorption at 495 and 580 nm) to FADH^−^, which is then reoxidised to FAD_ox_ without light [17,66,67]. Green light itself cannot photoreduce FAD_ox_ and start this redox cycle, but the magnetic orientation of birds under this spectrum is possible, as the results of various behaviour studies prove, but only for less than 1 hour. Birds lost their ability to use the magnetic compass in orientation experiments under green light condition after 1 hour of pre-exposure to total darkness/green light or 1 hour of testing in green light, in contrast to blue or turquoise light [47]. Our results of experiments in monochromatic green light with another peak wavelength (530 nm vs 565 nm) independently confirm this suggestion: under this light condition accumulated supply of FADH^·^ enables the second part of the redox cycle to run further to FADH^−^ and then to FAD_ox_ only if birds are tested immediately/shortly after the lights in an indoor aviary are turned off [31,38] or birds are kept in an outdoor aviary under full-spectrum natural light before tests [22,69]. The intensity of green light in our experiments was at least two times lower than in the majority of prior studies performed in different species ([35,42,45,47]; see Table S1 for details) and it can explain disorientation of pied flycatchers. However, the results of some other studies indicate that European robins and garden warblers were able to choose the appropriate migratory direction under the same or even lower intensity of green light (1 and 0.2-0.3 mW m^−2^; [22,36,39]).

In contradiction to our assumptions and the theory mentioned above, pied flycatchers were oriented under dim yellow light towards southwest in the natural magnetic field. The southwesterly direction corresponds to the expected migratory direction, which is typical for pied flycatchers from the eastern Baltic [53,64,70]. It should be noted that the same birds showed similar direction under different experimental conditions: full-spectrum natural light under stars/without them (Fig. 1 A, B) and low-intensity yellow light with access only to the magnetic field as a cue source (Fig. 2C). Along with highly expressed nocturnal restlessness before tests, these findings allow us to assume that our birds exhibited true migratory behaviour in all mentioned cases and use the magnetic field as a compass cue source in the absence of astronomical cues. Additionally, pied flycatchers predictably responded to the deflection of the horizontal component of the magnetic field (120° counterclockwise) and changed their orientation from southwest to northeast. All of this strongly indicates for the first time that pied flycatchers are able to choose migratory direction under dim yellow light using only the magnetic compass.

This finding is not in the line with orientation experiments in songbird migrants (garden warblers and European robins) conducted under 590 nm yellow (2.9 mW m^−2^ or 7 - 43×10^15^ quanta s^−1^ m^−2^) or 567.5 nm green-yellow (1 and 5 mW m^−2^ or 2.9 and 14×10^15^ quanta s^−1^ m^−2^) and resulted in disorientation [33,35,38,69]. There might be two explanations for these contradictive results:

1. Orientation of pied flycatchers under monochromatic yellow light is not true magnetic compass orientation but a ‘fixed direction’ response, previously shown in ‘darkness’ or dim red light with the same intensity (1 mW m^−2^; [44,45]). We might completely exclude the possibility that the birds showed such type of behaviour under dim yellow light in our autumn experiments only if they would be tested again in the magnetic field with an inverted vertical component or during spring migration and reverse their headings or show the seasonally appropriate migratory direction in spring, respectively. Both European robins and Australian silvereyes showed similar west-northwesterly direction regardless of the migratory season under different light conditions: total darkness, dim red light (1 mW m^−2^) and high-intensity green light (15 mW m^−2^ or 43×10^15^ quanta s^−1^ m^−2^; [34,36,40,44,45]). Therefore, the ‘fixed direction’ response is characterized by the same directional tendencies not only in spring and autumn (regardless of the migratory direction) but also in different species. However, orientation of pied flycatchers under dim yellow light differs from the typical ‘fixed direction’ response: the birds were oriented in appropriate southwesterly migratory direction (not towards the NW as other species do) and preferred different direction in darkness (Fig. 1C). Orientation of pied flycatchers in darkness seems to be a ‘fixed direction’ response: they were active and headed in the northeasterly direction which is not a seasonally appropriate migratory direction in autumn for this species [53,64,71]. This suggests that the southwesterly headings under dim yellow light may represent true magnetic compass orientation but not the type of response of the birds in darkness and dim red light.
2. As proposed in previous studies in newts and European robins [26,39], the yellow-light magnetic orientation in pied flycatchers can be explained by the existence of a low-sensitive long-wavelength (yellow-red) mechanism along with a high-sensitive short-wavelength (UV-blue) mechanism of light-dependent magnetoreception. Although the claim that European robins have a second peak of sensitivity to monochromatic light in the red spectrum [39] has not been confirmed in an independent study [45], support for the involvement of the long-wavelength mechanism in avian magnetoreception can be given by several behavioural studies. When yellow light was added to blue and green, the birds showed fixed directions towards the south under blue-and-yellow and towards the north under green-and-yellow in both spring and autumn [41]. According to the authors of this study, the cause of such unusual orientation might be a complex interaction of a short-wavelength and a long-wavelength receptors. In another study, European robins, pre-exposed to red light before tests, were oriented in a seasonal migratory direction under the same red light, in contrast to the birds pre-exposed to total darkness or full-spectrum light [42]. Results of this study indicate that the birds are able to orient under long-wavelength light but only after acclimation to specific light condition. A similar adaptation of the avian magnetic compass to new experimental condition was discovered in experiments with functional window: testing of European robins in the magnetic field weaker or stronger than the local one leads to disorientation [66]. However, after staying in the magnetic field with higher or lower intensity, birds are able to orient at the respective intensity [72,73]. Furthermore, it has been shown in neurophysiological and immunohistochemical studies that cells in the nucleus of the basal optic root and the optic tectum of homing pigeons exhibit responses to changes in the direction of the magnetic field with peaks under 503 and 582 nm light ([74], but see [51]) and illumination of 590 nm yellow light leads to a stronger antiserum labelling in Cry1a than under 565 nm green light [75].

## Conclusion

Many passerine birds are diurnal animals but migrate during the night developing nocturnal activity or restlessness (known as ‘Zugunruhe’; [76]) before the start of migration. However, the use of light-dependent magnetoreceptor at night faces challenges: dramatic changes in the spectral composition and intensity of light after local sunset. The spectral composition of light changes substantially during sunset and at twilight (civil, nautical, and astronomical) in the absence of artificial light pollution. After the disappearance of the sun’s disk under the horizon, red-shifted direct light and blue-shifted scattered light from the sun dominate during civil twilight [77] and then the spectrum shifts to the shorter wavelength, with an intensive blue peak [78] due to the absorption of long-wavelength light by the ozone layer (nautical twilight) [79]. At astronomical twilight, when the sun is more than 12° below the horizon and all of its signs disappear, the spectral composition depends on the presence or absence of moonlight and mostly long-wavelength. At full moon night the spectrum is close to the typical daylight spectrum ([80], see more details in [77]) and in the absence of the moon, starlight spectrum is long-wavelength-shifted with several peaks (at 560, 590, 630 and 684 nm [80,81]). In urban areas with light pollution, the spectral composition is similar to the light spectrum mentioned above at civil twilight, but it is strongly influenced by artificial light at nautical twilight and is fully dominated by such illumination with a broad peak centred at 590 nm starting with astronomical twilight and at night [78,80]. The intensity of light at visible wavelengths (400-700 nm) changes rapidly from 10^18^ quanta s^−1^ m^−2^ (photon quantity) or 1-10 W m^−2^ (irradiance) at sunset to 10^13^ quanta s^−1^ m^−2^ (a starlit night)/10^15^ quanta s^−1^ m^−2^ (a moonlit night) or 10^−4^ – 10^−5^ W m^−2^ and lower after the end of nautical twilight ([78,80,82]; Figure 3).

**Figure 3.**
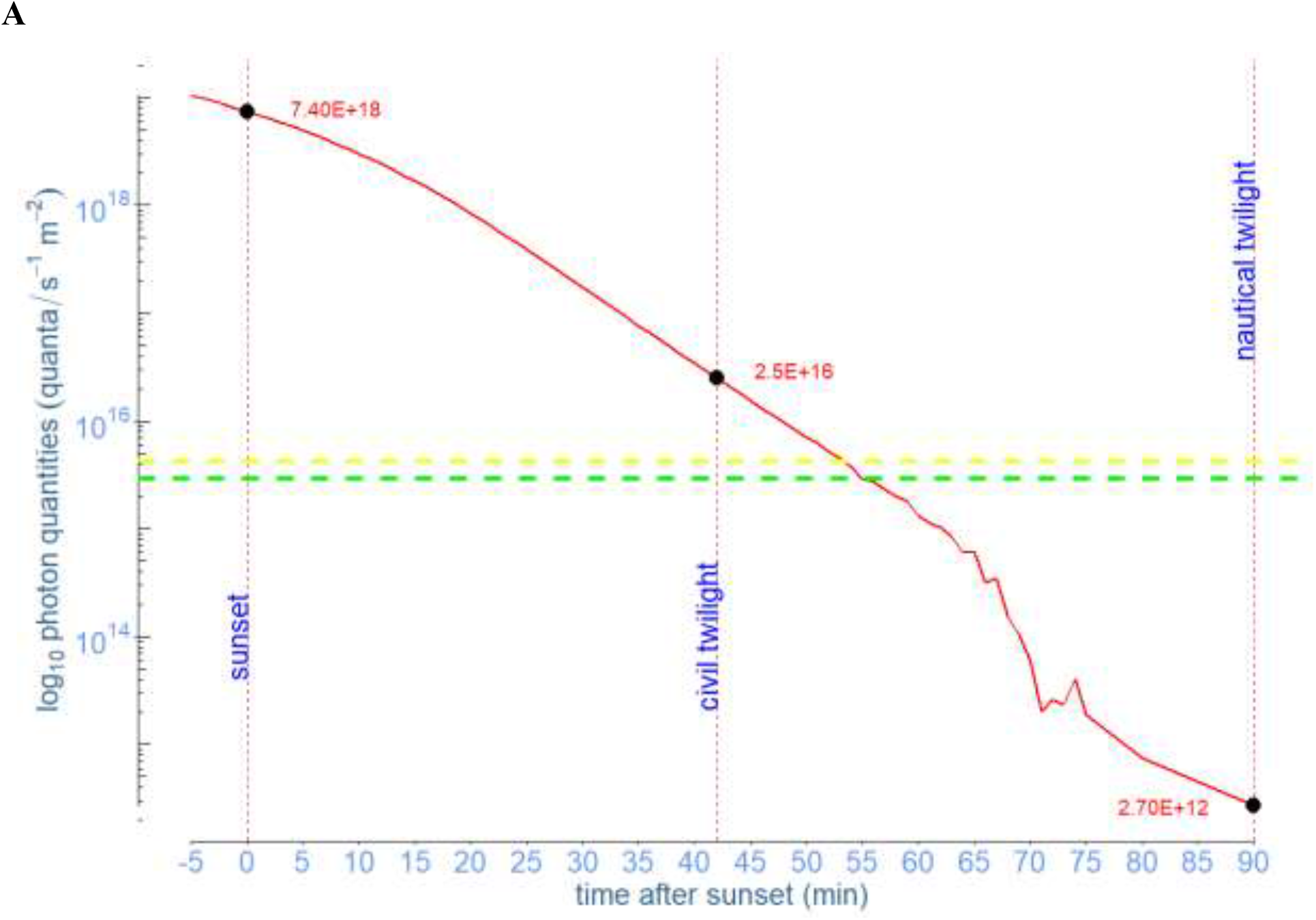
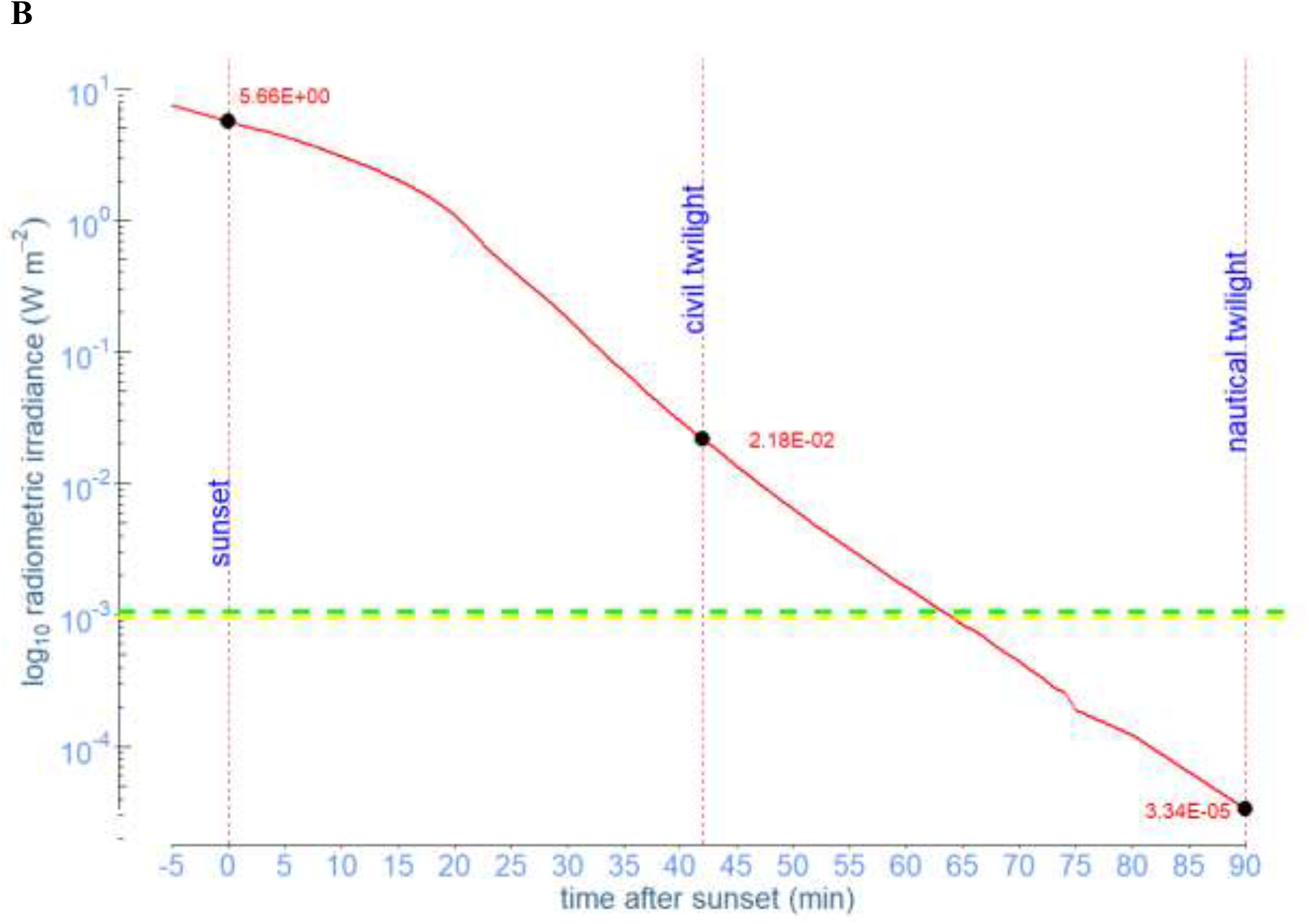
Intensity of natural light (photon quantity (A) and irradiance mW m^−2^ (B)) changes in downwelling illumination as a function of time after sunset measured at our experimental place. Green and yellow dashed lines indicate monochromatic light intensity (green and yellow) used in indoor experiments.

Currently, we do not know for sure when nocturnal migratory birds select their migratory direction: at sunset and civil twilight when they usually calibrate their compass systems [53,83,84] or show non-flight activity [85] or during the whole night from sunset to sunrise. Most species initiate their migratory flights in the wild or nocturnal restlessness in cages within 1–2 h after sunset, according to radiotelemetry, cage and radar studies [86–91]. Therefore, the avian magnetic compass should be adapted to different spectrum and light intensities that migratory birds face with throughout the night. And the existence of two light-dependent mechanisms which involve different cryptochromes might help them. One of the putative molecule candidates, Cry1a, is localized in the photoreceptor outer segments of the ultraviolet/violet (UV/V) cones in migratory (European robin and Eurasian blackcap) and non-migratory species (zebra finch and domestic chicken) and homing pigeon [92,93]. UV/V cones are the only cone type with transparent oil droplets (intracellular structures that usually play a role of spectral filters and cut off short-wavelength light) [94], so Cry1a, which has been suggested to be excited by blue light to form radical pairs ([95], but see [92]) is a good candidate for a high-sensitive short-wavelength receptor in this two receptors/mechanisms model. On the other hand, Cry4, which is currently considered to be the most likely light-dependent magnetoreceptor [96], has been detected in double cones and long-wavelength single (LWS) cones [96,97]. Avian LWS and double cones contain the oil droplets that cut off wavelengths below about 570 nm and 450 nm, respectively [94,98]. This light is required for cryptochrome photoreduction, thus Cry4 could play a role of a yellow-light sensor in the long-wavelength mechanism of avian magnetoreception.

## Acknowledgements

We are grateful to Julia Bojarinova and Dmitriy Sannikov for assistance in the experiments and analysis of Emlen funnels data. The authors would also like to express gratitude to Henrik Mouritsen for allowing us to use the magnetic coils belonging to his group, even though he had no role in designing and performing this study. We thank Nikita Chernetsov, Alexander Rotov and Luba Astakhova for their comments which greatly improved the earlier draft.

## Authors’ contributions

A.P. and N.R. designed research; A.P., N.R., F.C. and G.U. constructed the temporary wooden house and experimental equipment; A.P., N.R., A.F., A.Pr. and G.U. performed experiments, collected and analysed the data; A.P. and N.R. wrote the first draft of the manuscript. All authors commented on the manuscript and gave final approval for publication.

## Funding

Financial support for this study was made available by the Russian Foundation for Basic Research (grant 20-04-01059 to A.P.) and Zoological Institute RAS (research project 122031100261).

## Competing interests

The authors declare no competing interests.

## Data Availability

All data generated or analysed during this study are included in this manuscript (and its Supplementary Information files) or available from the corresponding author at reasonable request.

## Notes

### Competing Interest Statement

The authors have declared no competing interest.

### Summary of Updates

Abstract updated: one sentence was rewritten (see new version below). Our data contradict results of previous experiments under monochromatic yellow light and might indicate the previously proposed hypothesis of two different mechanisms in avian magnetoreception (a high-sensitive short-wavelength mechanism and a low-sensitive mechanism in the long-wavelength spectrum) has a right to exist. Results updated: we added results of experiments in darkness (see details below). Under darknes condition in 2022, the birds showed northestern direction (α = 39 deg, n = 10, r = 0.61, p = 0.02, Figure 1C) which does not correspond with autumn migratory direction typical for this species. Discussion updated: two sentences were rewritten (see new version below). 1) UV/V cones are the only cone type with transparent oil droplets (intracellular structures that usually play a role of spectral filters and cut off short-wavelength light)... 2) Avian LWS and double cones contain the oil droplets that cut off wavelengths below about 570 nm and 450 nm, respectively [94,98]. This light is required for cryptochrome photoreduction, thus Cry4 could play a role of a yellow-light sensor in the long-wavelength mechanism of avian magnetoreception. Main text updated (Materials and Methods, Results, Discussion): total darkness was changed to darkness

