## Supplementary materials for "Migratory birds are able to choose the appropriate migratory direction under dim yellow monochromatic light"

**Table S1**. Orientation of migratory birds in different light conditions.
p-value of the Rayleigh test: *P<0.05; **P<0.01; ***P<0.001; color indication in Results: **black** – normal orientation, **red** – disorientation/random, **blue** – unusual orientation response; OMF – oscillating magnetic field; mN – magnetic North; $8.7^{J}$ **– data from Johnsen et al., 2007**

| № | Species | Season | Colour/ experimental condition | Light source | Peak (nm) | Irradiance ($mW\ m^{-2}$) | Photon quantities ($*10^{15}$ quanta $s^{-1}\ m^{-2}$) | Result of orientation tests | Direction (deg) | Vector length | References |
|---|---|---|---|---|---|---|---|---|---|---|---|
| 1 | **Garden Warbler** *Sylvia borin* | autumn | White | Incandescent light bulb | - | - | 8.7 | **oriented** | 159 | $0.75^{***}$ | (Rappl et al., 2000) |
| | | | Blue | Glass filter, cool beam lamp | 443 | - | 8.7 | **oriented** | 166 | $0.55^{*}$ | |
| | | | Green | LED | 565 | - | 8.7 | **oriented** | 183 | $0.7^{**}$ | |
| | | | Orange-Yellow | LED | 584-592 (590) | - | 8.7 | **disoriented** | 236 | 0.34 | |
| | | | Red | LED | 626-635 (630) | - | 8.7 | **disoriented** | 5 | 0.14 | |
| | | | Green | LED | 527 | 0.25 | - | **oriented** | 216 | $0.38^{*}$ | (Bojarinova et al., 2020) |
| | | | Green / OMF applied locally to eye | LED | 527 | 0.25 | - | **oriented** | 189 | $0.49^{*}$ | |
| | | | Green / OMF | LED | 527 | 0.25 | - | **disoriented** | 106 | 0.11 | |
| 2 | **Australian silvereye** *Zosterops l. lateralis* | southern autumn | White | Tungsten light bulb | - | - | - | **oriented** | 34 | $0.63^{***}$ | (Wiltschko et al., 1993) |
| | | | Blue | Glass filter, cool beam | 443 | 2.7 | $8.7^{J}$ | **oriented** | 32 | $0.76^{***}$ | |

| | | | | | | | | | | | |
|---|---|---|---|---|---|---|---|---|---|---|---|
| | | | | lamp | | | | | | | |
| | | | Green | LED | 571 | 3 | 8.7[J] | **oriented** | 21 | 0.84*** | |
| | | | Red | LED | 633 | 3.9 | | **disoriented** | 182 | 0.08 | |
| | | | White | Tungsten light bulb (3.5 lux) | - | - | - | **oriented** | 18 | 0.91*** | (Munro et al., 1997) |
| | | | Green | LED | 571 | 3 | - | **oriented** | 14 | 0.94*** | |
| | | | Red | LED | 633 | 3.9 | - | **disoriented** | 40 | 0.1 | |
| | | | Green | LED | 565 | 0.2 | - | **oriented /less activity** | 354 | 0.64* | (Wiltschko et al., 2000a) |
| | | | Green | LED | 565 | 2.1 | 6.5[J] | **oriented** | 12 | 0.93*** | |
| | | | Green | LED | 565 | 7.5 | 22 | **oriented** | 9 | 0.95*** | |
| | | | Green | LED | 565 | 15 | 44 | **oriented /shifted response** | 304 | 0.88*** | |
| | | southern spring | Green | LED | 565 | 2.1 | | **oriented** | 187 | 0.91*** | (Wiltschko et al., 2000b) |
| | | | Green | LED | 565 | 7.5 | 22 | **oriented** | 200 | 0.91*** | |
| | | | Green | LED | 565 | 15 | 44 | **oriented /shifted response** | 304 | 0.84*** | |
| | | | Blue | LED | 424 | 2.8 | 7 | **oriented** | 192 | 0.95*** | (Wiltschko et al., 2003) |
| | | | Blue / vert. comp. inverted | LED | 424 | 2.8 | 7 | **oriented** | 355 | 0.84*** | |
| | | | Blue | LED | 424 | 20 | 43 | **oriented /axial response** | 93–273 | 0.56* | |
| | | | Green | LED | 565 | 2.1 | 7 | **oriented** | 188 | 0.87*** | |
| | | | Green / vert. comp. inverted | LED | 565 | 2.1 | 7 | **oriented** | 360 | 0.74*** | |
| | | | Green | LED | 565 | 15 | 43 | **oriented / tendency to** | 283 | 0.85*** | |

| | | | | | | | | | | | |
|---|---|---|---|---|---|---|---|---|---|---|---|
| | | | | | | | | **NW** | | | |
| | | | Green / vert. comp. inverted | LED | 565 | 15 | 43 | **oriented /tendency to NW** | 293 | 0.63** | |
| | | | Green/ mN=240˚ | LED | 565 | 15 | 43 | **oriented/ shifted response to SE** | 143 | 0.92*** | |
| | | | Green | LED | 565 | 2 | - | **oriented** | 175 | 0.86*** | (Wiltschko et al., 2008) |
| | | | Red | LED | 645 | 1 | - | **oriented/ tendency to W** | 276 | 0.89*** | |
| | | | Red | LED | 645 | 2 | - | **disoriented** | 303 | 0.48 | |
| | | | Red/vert. comp. inverted | LED | 645 | 1 | - | **oriented/ tendency to W** | 284 | 0.92*** | |
| | | | Red/mN=270˚ | LED | 645 | 1 | - | **oriented/ shifted response** | 197 | 0.92*** | |
| | | | Green | LED | 565 | 2 | - | **oriented** | 194 | 0.84*** | |
| | | | Green/beak anaesthetized | LED | 565 | 2 | - | **oriented** | 190 | 0.73** | |
| | | | Red | LED | 645 | 1 | - | **oriented /tendency to W** | 280 | 0.61*** | |
| | | | Red/beak anaesthetized | LED | 645 | 1 | - | **disoriented** | 98 | 0.06 | |
| | | | Total darkness | | | | - | **oriented** | 245 | 0.77** | |
| | | | Green | LED | 565 | 1.9 | - | **oriented** | 176 | 0.95*** | (Wiltschko et al., 2014a) |
| | | | UV | LED | 373 | 0.3 | - | **oriented** | 178 | 0.76*** | |
| | | | UV | LED | 373 | 2.8 | - | **oriented /shifted** | 59 | 0.94*** | |

| | | | | | | | | | | |
|---|---|---|---|---|---|---|---|---|---|---|
| | | | | | | | | **response** | | |
| | | | UV/vert. comp. inverted | LED | 373 | 2.8 | - | **oriented/ shifted response** | 130 | $0.95^{***}$ |
| 3 | European robin *Erithacus rubecula* | spring | White | Tungsten light bulb | | 3.7 | - | **oriented** | 24 | $0.78^{***}$ | (Wiltschko and Wiltschko, 1995) |
| | | | Green | LED | 571 | 3 | - | **oriented** | 13 | $0.89^{***}$ | |
| | | | Red | LED | 633 | 3.9 | - | **disoriented** | 162 | 0.32 | |
| | | | Blue | LED | 443 | 3.9 | 8.7 | **oriented** | 13 | $0.72^{***}$ | (Wiltschko and Wiltschko, 1999) |
| | | | Green | LED | 565 | 3 | 8.7 | **oriented** | 360 | $0.9^{***}$ | |
| | | | Green/mN=120˚ | LED | 565 | 3 | 8.7 | **oriented /shifted response** | 131 | $0.84^{***}$ | |
| | | | Orange-Yellow | LED | 584-592 (590) | 2.9 | 8.7 | **disoriented** | 229 | 0.4 | |
| | | | Blue | Glass filter, cool beam lamp | 424 | - | 7 | **oriented** | 6 | $0.76^{***}$ | (Wiltschko and Wiltschko, 2001) |
| | | | Blue | Glass filter, cool beam lamp | 424 | - | 43 | **oriented /axial response** | 272-92 | $0.61^{*}$ | |
| | | | Turquoise | LED | 510 | - | 7 | **oriented** | 24 | $0.85^{***}$ | |
| | | | Turquoise | LED | 510 | - | 43 | **oriented/ shifted response** | 348 | $0.62^{***}$ | |
| | | | Green | LED | 565 | - | 7 | **oriented** | 13 | $0.91^{***}$ | |
| | | | Green | LED | 565 | - | 43 | **oriented/ axial response** | 271-91 | $0.78^{**}$ | |
| | | | Yellow | LED | 590 | - | 7 | **disoriented** | 247 | 0.05 | |
| | | | Yellow | LED | 590 | - | 43 | **disoriented** | 17 | 0.29 | |

| | | | | | | | | | | | |
|---|---|---|---|---|---|---|---|---|---|---|---|
| | | | White | incandescent light bulb | | 24.4 | - | **oriented** | 5 | 0.96*** | |
| | | | Red | LED | 645 | 2.1 | 6-7 | **disoriented** | 46 | 0.34 | |
| | | | Red | LED | 645 | 13 | 43 | **disoriented** | 37 | 0.34 | |
| | | | Red /1 h red pre-exposure | LED | 645 | 2.1 | 6-7 | **oriented** | 25 | 0.81*** | |
| | | | Red /1 h red pre-exposure | LED | 645 | 13 | 43 | **oriented** | 32 | 0.62*** | |
| | | | Green /1 h red pre-exposure | LED | 565 | 2.4 | 6-7 | **oriented** | 357 | 0.56* | |
| | | | Red | LED | 645 | 1.8 | 6-7 | **disoriented** | 278 | 0.3 | |
| | | | Green | LED | 565 | 2.1 | 6-7 | **oriented** | 6 | 0.94*** | |
| | | | Red /1 h red pre-exposure | LED | 645 | 1.8 | 6-7 | **oriented** | 358 | 0.79*** | |
| | | | Green /1 h red pre-exposure | LED | 565 | 2.1 | 6-7 | **disoriented** | 357 | 0.49 | |
| | | | Red /1 h dark pre-exposure | LED | 645 | 1.8 | 6-7 | **disoriented** | 322 | 0.24 | |
| | | | Green /1 h dark pre-exposure | LED | 565 | 2.1 | 6-7 | **oriented** | 17 | 0.88*** | (Wiltschko et al., 2004a) |
| | | | Green | LED | 565 | 2.1 | 7 | **oriented** | 17 | 0.93*** | |
| | | | Green - Yellow | LED | 565 and 590 | 2.1 (G), 2.0 (Y) | 14 | **oriented** | 25 | 0.52* | |
| | | | Blue - Yellow | LED | 424 and 590 | 2.8 (B), 2.0 (Y) | 14 | **oriented /shifted response** | 200 | 0.81*** | (Wiltschko et al., 2004b) |
| | | | Blue | LED | 424 | - | 8 | **oriented** | 8 | 0.76*** | (Wiltschko et al., 2007) |
| | | | Blue | LED | 424 | - | 36 | **oriented /axial response** | 3-183 | 0.69** | |
| | | | Blue | LED | 424 | - | 54 | **oriented /axial** | 19-199 | 0.39 | |

| | | | | | | | | | | | |
|---|---|---|---|---|---|---|---|---|---|---|---|
| | | | | | | | | **response** | | | |
| | | | Blue | LED | 424 | - | 72 | **oriented /axial response** | 6-186 | 0.6$^{**}$ | |
| | | | Turquoise | LED | 502 | - | 8 | **oriented** | 5 | 0.96$^{***}$ | |
| | | | Turquoise | LED | 502 | - | 36 | **oriented /axial response** | 98-278 | 0.61$^{**}$ | |
| | | | Turquoise | LED | 502 | - | 54 | **oriented** | 5 | 0.89$^{***}$ | |
| | | | Turquoise | LED | 502 | - | 72 | **oriented** | 13 | 0.93$^{***}$ | |
| | | | Green | LED | 565 | - | 8 | **oriented** | 9 | 0.91$^{***}$ | |
| | | | Green | LED | 565 | - | 36 | **disoriented** | 55-235 | 0.27 | |
| | | | Green | LED | 565 | - | 54 | **oriented /axial response** | 102-282 | 0.7$^{**}$ | |
| | | | Green | LED | 565 | - | 72 | **oriented /axial response** | 19-199 | 0.68$^{**}$ | |
| | | | UV | LED | 373 | - | 8 | **oriented /axial response** | 89-269 | 0.5$^{*}$ | |
| | | | UV | LED | 373 | - | 0.8 | **oriented** | 8 | 0.96$^{***}$ | |
| | | | Green | LED | 565 | 2 | - | **oriented** | 14 | 0.88$^{***}$ | (Wiltschko et al., 2008) |
| | | | Red | LED | 645 | 1 | - | **oriented/ tendency to W** | 273 | 0.85$^{***}$ | |
| | | | Total darkness | | | 0 | - | **oriented / tendency to W** | 278 | 0.88$^{***}$ | |
| | | | Green | LED | 565 | 2 | - | **oriented** | 9 | 0.86$^{***}$ | (Stapput et al., 2008) |
| | | | Total darkness | | | - | - | **oriented / tendency to NW** | 300 | 0.81$^{***}$ | |

| | | | | | | | | | | | |
|---|---|---|---|---|---|---|---|---|---|---|---|
| | | | Total darkness/ vert. comp. inverted | | | - | - | **oriented / tendency to NW** | 302 | 0.81*** | |
| | | | Total darkness/ mN=180˚ | | | - | - | **oriented / tendency to SE** | 125 | 0.94*** | |
| | | | Total darkness/ horiz. comp. inverted + OMF | | | - | - | **oriented / tendency to W** | 278 | 0.65** | |
| | | | Total darkness/ beak anesthetized | | | - | - | **disoriented** | 73 | 0.21 | |
| | | | White | 15 W light bulb | - | - | - | **oriented** | 6 | 0.85*** | (Wiltschko et al., 2014a) |
| | | | Green | LED | 565 | 1.9 | - | **oriented** | 5 | 0.96*** | |
| | | | Turquoise | LED | 502 | 2.2 | - | **oriented** | 12 | 0.98*** | |
| | | | UV | LED | 373 | 0.3 | 0.8 | **oriented** | 13 | 0.96*** | |
| | | | UV | LED | 373 | 2.8 | 8 | **oriented/ axial response** | 86-266 | 0.44** | |
| | | | UV | LED | 373 | 5.7 | - | **oriented/ axial response** | 13-193 | 0.76*** | |
| | | | UV/ vert.comp. inverted | LED | 373 | 0.3 | - | **oriented/ shifted response** | 173 | 0.71*** | |
| | | | UV/ OMF added | LED | 373 | 0.3 | - | **disoriented** | 174 | 0.15 | |
| | | | UV/ beak anesthetized | LED | 373 | 0.3 | - | **oriented** | 20 | 0.97*** | |
| | | | UV+Yellow | LED | 373 and 585 | 0.3 (UV), 1.7 (Y) | - | **weakly oriented** | 55 | 0.57* | |
| | | | UV+Yellow | LED | 373 | 2.8 (UV), | - | **oriented/** | 91 | 0.76*** | |

| | | | | | | | | | | | |
|---|---|---|---|---|---|---|---|---|---|---|---|
| | | | | | and 585 | 1.7 (Y) | | **tendency to E** | | | |
| | | | UV+Yellow/ OMF added | LED | 373 and 585 | 2.8 (UV), 1.7 (Y) | - | **oriented/ tendency to SE** | 128 | $0.53^{*}$ | |
| | | | UV+Yellow/ beak anesthetized | LED | 373 and 585 | 2.8 (UV), 1.7 (Y) | - | **disoriented** | 46 | 0.11 | |
| | | | Green/white light pre-exposure | LED | 564 | 1.9 | 8 | **oriented** | 353 | $0.6^{*}$ | (Wiltschko et al., 2014b) |
| | | | Blue/1 h dark pre-exposure | LED | 424 | 2.4 | 8 | **oriented** | 11 | $0.99^{***}$ | |
| | | | Turquoise/1 h dark pre-exposure | LED | 502 | 2.11 | 8 | **oriented** | 354 | $0.96^{***}$ | |
| | | | Green/1 h dark pre-exposure | LED | 565 | 1.9 | 8 | **oriented/ axial response** | 77-257 | $0.6^{*}$ | |
| | | | Blue/1 h blue light pre-exposure | LED | 424 | 2.4 | 8 | **oriented/ axial response** | 16-196 | $0.72^{***}$ | |
| | | | Turquoise/1 h turquoise light pre-exposure | LED | 502 | 2.11 | 8 | **oriented** | 6 | $0.95^{***}$ | |
| | | | Green/1 h green light pre-exposure | LED | 565 | 1.9 | 8 | **disoriented** | 325 | 0.35 | |
| | | autumn | White | 1000W ozone-free xenon arc lamp | | 1 | 3.9 | **disoriented** | 244-64 | 0.13 | (Muheim et al., 2002) |
| | | | White | 1000W | | 10 | 39 | **disoriented** | 41-221 | 0.34 | |

| | | | | | | | | | | |
|---|---|---|---|---|---|---|---|---|---|---|
| | | | | ozone-free xenon arc lamp | | | | | | |
| | | | White (before tests) | 1000W ozone-free xenon arc lamp | | 20 | 79 | **oriented/ tendency to W** | 282 | 0.34* |
| | | | White (after tests) | 1000W ozone-free xenon arc lamp | | 20 | 79 | **disoriented** | 229 | 0.3 |
| | | | Green | 1000W ozone-free xenon arc lamp + colour filter | 560.5 | 1 | 2.9 | oriented | 215 | 0.49** |
| | | | Green | 1000W ozone-free xenon arc lamp + colour filter | 560.5 | 5 | 14 | **oriented/ axial response** | 214-34 | 0.41* |
| | | | Green | 1000W ozone-free xenon arc lamp+ colour filter | 560.5 | 10 | 29 | **disoriented** | 216-36 | 0.4 |
| | | | Green-Yellow | 1000W ozone-free xenon arc lamp+ colour filter | 567.5 | 1 | 2.9 | **disoriented** | 74 | 0.23 |
| | | | Green-Yellow | 1000W ozone-free | 567.5 | 5 | 14 | **disoriented** | 177 | 0.29 |

| | | | | | | | | | | | |
|---|---|---|---|---|---|---|---|---|---|---|---|
| | | | | xenon arc lamp+ colour filter | | | | | | | |
| | | | Green-Yellow | 1000W ozone-free xenon arc lamp+ colour filter | 567.5 | 10 | 29 | **disoriented** | 238 | 0.3 | |
| | | | Red | 1000W ozone-free xenon arc lamp+ colour filter | 617 | 1 | 3.2 | **oriented/ tendency to W** | 277 | 0.71*** | |
| | | | Red | 1000W ozone-free xenon arc lamp+ colour filter | 617 | 5 | 16 | **oriented/ tendency to SW** | 259 | 0.48* | |
| | | | Red | 1000W ozone-free xenon arc lamp+ colour filter | 617 | 10 | 32 | **disoriented** | 315 | 0.34 | |
| | | | White | incandescent light bulb | | 24.4 | | **oriented** | 201 | 0.64*** | (Wiltschko et al., 2004a) |
| | | | Green | LED | 565 | 2.1 | 6-7 | **oriented** | 200 | 0.68*** | |
| | | | Red/1 h dark pre-exposure | | 645 | 1.6 | 6-7 | **oriented** | 181 | 0.62** | |
| | | | Green | LED | 565 | 2.1 | 7 | **oriented** | 200 | 0.68*** | (Wiltschko et al., 2004b) |
| | | | Green + Yellow | LED | 565 and 590 | 2.1 (G), 2.0 (Y) | 14 | **oriented/ shifted response** | 22 | 0.63*** | |
| | | | Blue + Yellow | LED | 424 | 2.8 (B), | 14 | **oriented/** | 132 | 0.52* | |

| | | | | | | | | | | | |
|---|---|---|---|---|---|---|---|---|---|---|---|
| | | | | | and 590 | 2.0 (Y) | | **shifted response** | | | |
| | | | Green | LED | 565 | 2 | - | **oriented** | 190 | $0.73^{***}$ | (Wiltschko et al., 2008) |
| | | | Red | LED | 645 | 1 | - | **oriented/ tendency to NW** | 289 | $0.72^{***}$ | |
| | | | Total darkness | | | | - | **oriented/ tendency to NW** | 303 | $0.74^{***}$ | |
| | | | White | 15 W light bulb | - | - | - | **oriented** | 176 | $0.85^{***}$ | (Wiltschko et al., 2014a) |
| | | | UV | LED | 373 | 0.3 | - | **oriented** | 182 | $0.53^{***}$ | |
| | | | UV | LED | 373 | 2.8 | - | **oriented/ axial response** | 78-258 | $0.63^{***}$ | |

**Table S2. Raw data of all orientation tests.**

NA – not active, NS – not significant; magnetic North is 0; **280** – only one significant result or difference between two tests is >120

***Orientation of pied flycatchers in full-spectrum light condition during autumn migration 2021.***

| | | Stars, outdoors 29.08.2021-08.09.2021 | | | | | | Simulated overcast (under plexiglass), outdoors 08.09.2021-20.09.2021 | | | | |
|---|---|---|---|---|---|---|---|---|---|---|---|---|
| **№** | **Ring** | **Test 1** | **Test 2** | **Test 3** | **Test 4** | **Test 5** | **Mean** | **Test 1** | **Test 2** | **Test 3** | **Test 4** | **Mean** |
| 1 | VC72576 | NS | **243** | **140** | **220** | **123** | **181** | NS | NS | **120** | **110** | **115** |
| 2 | VC71793 | **186** | **100** | **230** | **255** | | **202** | NA | **253** | NS | **100** | **177** |
| 3 | VC71839 | NS | **28** | NA | NS | | **28** | NS | NA | NA | NA | |
| 4 | VC71842 | NS | **78** | **190** | **183** | | **157** | NA | NA | NS | NA | |
| 5 | VC72624 | NS | NS | **165** | **240** | | **203** | **248** | **143** | **238** | NS | **215** |
| 6 | VC71917 | NS | **23** | **175** | **238** | **218** | **215** | **180** | NA | | | **180** |
| 7 | VC71939 | NS | **215** | NS | **170** | | **193** | **248** | **120** | NA | **120** | **150** |
| 8 | VC71938 | **320** | NS | **213** | | | **267** | NS | NS | **326** | **245** | **286** |
| 9 | VC71950 | NS | **180** | **238** | **105** | | **176** | **168** | **123** | **163** | **310** | **163** |
| 10 | VC71951 | **135** | **293** | **135** | | | **154** | NA | NA | NS | **105** | **105** |
| 11 | VC72685 | **8** | NS | **208** | **220** | **185** | **213** | **195** | **80** | | | **138** |
| 12 | VC72687 | NS | **245** | **325** | **303** | | **292** | NS | **233** | | | **233** |
| 13 | VC71961 | NS | NS | | | | | | | | | |
| 14 | VC72774 | **238** | **158** | NS | | | **198** | NS | **155** | **253** | NS | **204** |
| 15 | VC72781 | **243** | **240** | **235** | **233** | | **238** | **100** | **260** | | | **180** |
| 16 | VC72799 | **170** | **165** | **183** | **205** | | **181** | **158** | **180** | NS | **250** | **194** |
| 17 | VC72871 | NS | NS | **213** | **NS** | | **213** | **290** | **200** | **140** | NS | **205** |
| 18 | VC76027 | NS | **298** | **183** | **293** | | **266** | NS | NS | NS | NS | |
| 19 | VC76134 | **118** | **280** | **330** | **310** | | **312** | NS | NS | | | |
| 20 | VC76315 | | | | | | | **160** | **280** | | | **220** |

*Orientation of pied flycatchers in green light condition during autumn migration 2021.*

| № | Ring | NMF, indoors 24.08.2021-21.09.2021 | | | | | 120° CCW CMF, indoors 24.08.2021-21.09.2021 | | | | |
|---|---|---|---|---|---|---|---|---|---|---|---|
| | | Test 1 | Test 2 | Test 3 | Test 4 | Mean | Test 1 | Test 2 | Test 3 | Test 4 | Mean |
| 1 | VC72576 | 100 | 100 | 80 | 295 | 83 | 150 | 202 | 143 | 245 | 183 |
| 2 | VC71793 | NS | 290 | NS | NS | 290 | NS | NS | 245 | NS | 245 |
| 3 | VC71839 | NS | NS | NS | NS | | NS | NS | NS | NS | |
| 4 | VC71842 | NS | NS | NS | 23 | 23 | 213 | NS | NS | | 213 |
| 5 | VC72624 | NS | 135 | 215 | NS | 175 | NS | 30 | NS | NS | 30 |
| 6 | VC71917 | 30 | 342 | 210 | 88 | 35 | NS | NS | NS | 128 | 128 |
| 7 | VC71939 | 133 | 55 | NS | NS | 94 | 288 | 80 | NS | | 4 |
| 8 | VC71938 | NS | NS | NS | | | 145 | NS | NS | | 145 |
| 9 | VC71950 | NS | NS | NS | | | 290 | NS | NS | | 290 |
| 10 | VC71951 | 190 | 285 | NS | | 238 | NA | NS | NS | | |
| 11 | VC72685 | NS | NS | NS | | | 15 | NS | NS | 295 | 335 |
| 12 | VC72687 | NS | NS | NS | | | NS | NS | NS | | |
| 13 | VC71961 | NS | NS | | | | NS | NS | NS | | |
| 14 | VC72774 | NS | 260 | | | 260 | NS | NS | | | |
| 15 | VC72781 | NS | NS | | | | 80 | NS | | | 80 |
| 16 | VC72799 | 130 | NS | | | 130 | NS | NS | | | |
| 17 | VC72871 | NS | NS | | | | NS | NS | | | |
| 18 | VC76027 | NS | NS | | | | NS | 270 | | | 270 |
| 19 | VC76134 | NS | 130 | | | 130 | NS | 230 | | | 230 |
| 20 | VC76315 | NS | NS | | | | NS | | | | |

*Orientation of pied flycatchers in yellow light condition during autumn migration 2021.*

| | | NMF, indoors 31.08.2021-22.09.2021 | | | | | | 120° CCW CMF, indoors 31.08.2021-22.09.2021 | | | | | |
|---|---|---|---|---|---|---|---|---|---|---|---|---|---|
| № | Ring | Test 1 | Test 2 | Test 3 | Test 4 | Test 5 | Mean | Test 1 | Test 2 | Test 3 | Test 4 | Test 5 | Mean |
| 1 | VC72576 | **325** | NS | **270** | **290** | **125** | **289** | **23** | **290** | **310** | | | **325** |
| 2 | VC71793 | **153** | NS | NA | **125** | | **139** | **93** | **190** | **120** | | | **132** |
| 3 | VC71839 | **180** | NS | **200** | | | **190** | **173** | NS | NS | **130** | **NS** | **152** |
| 4 | VC71842 | NS | NS | NS | NS | | | NS | NA | NS | NS | | |
| 5 | VC72624 | NS | NS | NS | NS | | | NS | NS | NS | NS | | |
| 6 | VC71917 | NS | **228** | **130** | NA | | **179** | NS | NA | **67** | **280** | | **354** |
| 7 | VC71939 | NS | NS | **235** | **320** | | **278** | NA | **NS** | **NA** | | | |
| 8 | VC71938 | NS | NS | NS | NS | | | **33** | NS | NS | NS | | **33** |
| 9 | VC71950 | NS | **205** | **275** | **120** | | **203** | **208** | **165** | **150** | **20** | NS | **160** |
| 10 | VC71951 | NS | **313** | **210** | NS | | **262** | NS | **65** | **155** | **130** | | **118** |
| 11 | VC72685 | NS | NS | **125** | **105** | | **115** | NS | NS | **130** | **NA** | | **130** |
| 12 | VC72687 | NS | NS | NS | **273** | | **273** | NS | NS | **20** | **NS** | | **20** |
| 13 | VC71961 | NS | NS | NS | NS | | | NS | NS | NS | | | |
| 14 | VC72774 | NS | NS | **70** | **210** | | **140** | NS | NS | NS | **350** | | **350** |
| 15 | VC72781 | NS | **180** | **275** | NS | | **228** | NS | **53** | NS | | | **53** |
| 16 | VC72799 | **168** | NS | NS | | | **168** | **248** | NS | **250** | **100** | **300** | **257** |
| 17 | VC72871 | **150** | NS | **170** | | | **160** | **335** | NS | **265** | **80** | | **336** |
| 18 | VC76027 | NS | NS | NS | NS | | | NS | NS | **115** | | | **115** |
| 19 | VC76134 | NS | NS | | | | | **10** | **NS** | | | | **10** |
| 20 | VC76315 | **240** | NA | | | | **240** | NA | **40** | | | | **40** |

*Orientation of pied flycatchers in full-spectrum light condition during autumn migration 2022.*

| № | Ring | Simulated overcast (under plexiglass), outdoors 24.08.2022-03.09.2022 | | | | | | | | "Darkness", indoors 17.09.2022-21.09.2022 | | | | |
|---|---|---|---|---|---|---|---|---|---|---|---|---|---|---|
| | | Test 1 | Test 2 | Test 3 | Test 4 | Test 5 | Test 6 | Test 7 | Mean | Test 1 | Test 2 | Test 3 | Test 4 | Mean |
| 1 | 134 | 190 | 310 | 43 | 285 | 283 | 20 | 220 | 293 | 70 | NA | | | 70 |
| 2 | 133 | NS | NS | NS | 250 | 15 | 170 | | 235 | NS | NS | NS | NS | |
| 3 | 114 | 230 | 330 | NS | 243 | | | | 265 | 355 | 30 | | | 12 |
| 4 | 420 | NA | 245 | 245 | NS | 270 | | | 253 | 75 | 145 | | | 110 |
| 5 | 121 | NS | 220 | NS | 335 | 273 | | | 275 | NS | NS | 25 | 340 | 3 |
| 6 | 124 | NS | NS | 175 | 215 | 63 | 198 | | 178 | NS | NS | 298 | 37 | 348 |
| 7 | 427 | NS | 335 | 295 | 5 | 308 | 270 | | 314 | NS | NS | NA | NS | |
| 8 | 152 | 315 | 25 | NS | 0 | 85 | | | 15 | 105 | 100 | | | 103 |
| 9 | 181 | NS | NS | 270 | NS | 300 | | | 285 | 250 | 350 | | | 300 |
| 10 | 165 | NS | 183 | 75 | NS | | | | 129 | | | | | |
| 11 | 166 | 90 | 270 | NS | 90 | 305 | 340 | | 354 | NS | 60 | | | 60 |
| 12 | 476 | 230 | 195 | 270 | NS | | | | 232 | NS | 0 | | | 0 |
| 13 | 170 | NS | NS | 223 | 55 | 215 | | | 204 | 200 | 95 | NS | 10 | 93 |

*Orientation of pied flycatchers in yellow light condition during autumn migration 2022.*

| № | Ring | NMF, indoors 06.09.2022-18.09.2022 | | | | | | | | 120° CCW CMF, indoors 06.09.2022-18.09.2022 | | | | | | | |
|---|---|---|---|---|---|---|---|---|---|---|---|---|---|---|---|---|---|
| | | Test 1 | Test 2 | Test 3 | Test 4 | Test 5 | Test 6 | Test 7 | Mean | Test 1 | Test 2 | Test 3 | Test 4 | Test 5 | Test 6 | Test 7 | Mean |
| 1 | 134 | NS | 217 | NA | 210 | | | | 214 | 56 | NA | NS | NS | | | | 56 |
| 2 | 133 | 145 | NS | NS | NS | 260 | NS | | 203 | 293 | 35 | 200 | NS | | | | 291 |
| 3 | 114 | NS | NS | NA | 199 | NA | NA | 150 | 175 | NA | NA | NA | 100 | NS | NS | | 100 |
| 4 | 420 | NS | NS | 150 | 145 | NS | | | 148 | 350 | 115 | NS | 130 | | | | 93 |
| 5 | 121 | NS | 90 | NS | NS | 105 | 115 | 128 | 110 | 335 | 330 | NS | NS | | | | 333 |
| 6 | 124 | NA | 186 | NA | NA | | | | 186 | NA | NA | NA | 85 | NA | NA | | 85 |
| 7 | 427 | NA | NS | NA | NS | | | | | 125 | NA | NS | 315 | NA | | | 40 |
| 8 | 152 | 164 | 225 | NS | NS | NS | 285 | | 225 | 14 | 28 | NS | 140 | | | | 51 |
| 9 | 181 | NS | 250 | 100 | NS | 170 | NS | | 172 | 55 | NS | NS | 135 | | | | 95 |
| 10 | 165 | 150 | 220 | NS | 223 | | | | 199 | | | | | | | | |
| 11 | 166 | NS | NS | NA | 176 | NS | NS | | 176 | 100 | 340 | NA | NA | | | | 40 |
| 12 | 476 | 200 | 150 | 220 | NA | NA | NS | | 191 | NA | NA | 345 | 130 | NA | NA | | 58 |
| 13 | 170 | 245 | NA | NS | NS | | | | 245 | NS | 120 | NS | 75 | | | | 98 |

**Table S3. The results of maximum likelihood estimation.**

The best fitted models are listed in corresponding column (model names are from Fitak & Johnsen, 2017)

| **Light conditions** | **Field** | **models with ΔAIC < 2** | **Comment** | **Estimated mean(s)** |
|---|---|---|---|---|
| Darkness | NMF | M5A, M2A | Bimodal, with strongly supported unimodal alternative | 85° and 349° for bimodal alternative, 40° for unimodal |
| Outdoor with stars | NMF | M2A, M2B | Unimodal | 212° |
| Outdoor under plexiglas | NMF | M2A | Unimodal | 211° |
| Monochromatic green | NMF | M1 | Uniform | |
| Monochromatic green | CMF | M1 | Uniform | |
| Monochromatic yellow | NMF | M2A | Unimodal | 195° |
| Monochromatic yellow | CMF | M2A | Unimodal | 56° |

**Figure S1. The temporary experimental chamber outside (A) and inside (a 3D model; B).**

1 – an aluminium frame with green and yellow LEDs strips; 2 – the magnetic coils; 3 – a Plexiglass lid on the top of Emlen funnel (a diffusor, 70% light transmission); 3– Emlen funnel with a tested bird; 4 – a temporary constructed wooden house.

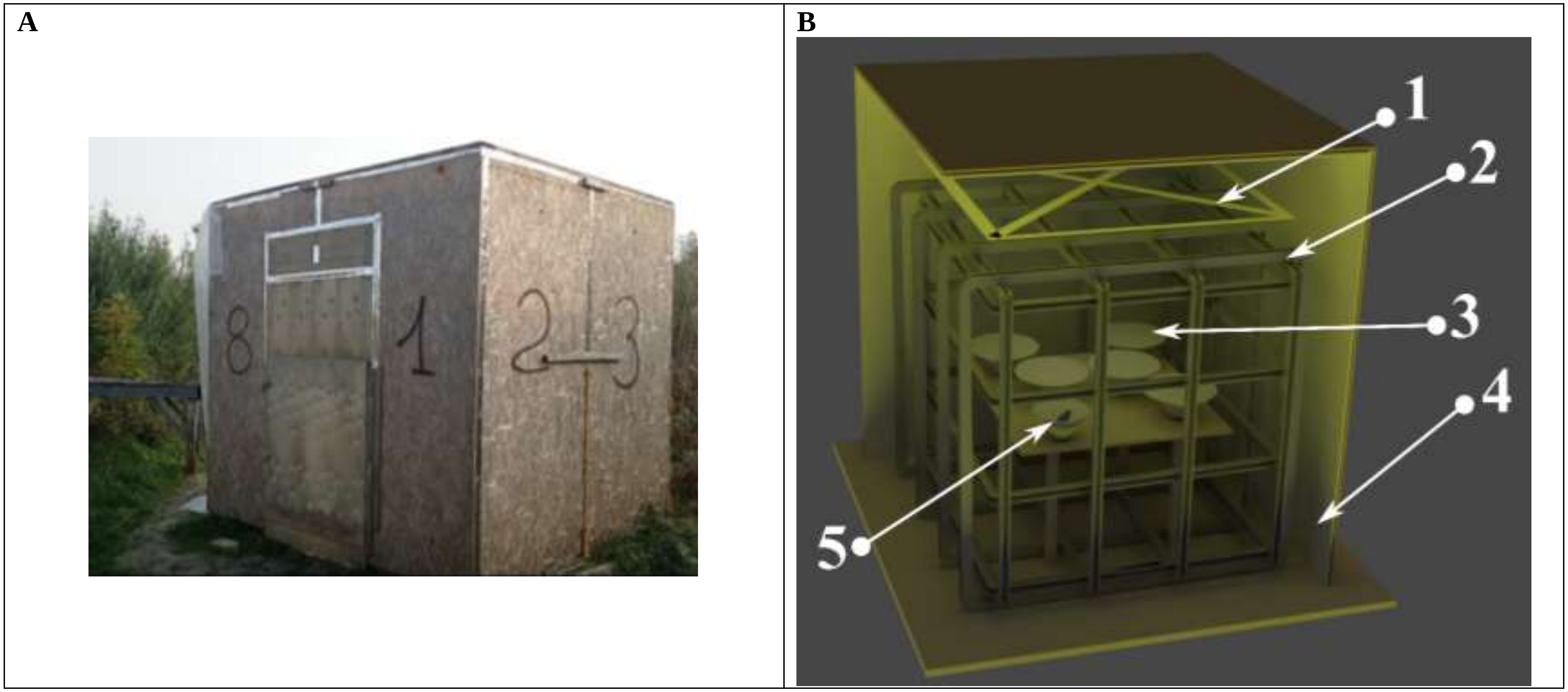

**Figure S2. Spectrometric measurements of monochromatic LEDs light used in indoor experiments (in the laboratory chamber).**

The green and yellow lines indicate the spectrum of the monochromatic light (green and yellow, consequently) inside the Emlen funnel under Plexiglass lids. All spectra are standardized by taking the maximum value within the measured wavelength interval as 1.0, similar to Muheim et al., 2002.

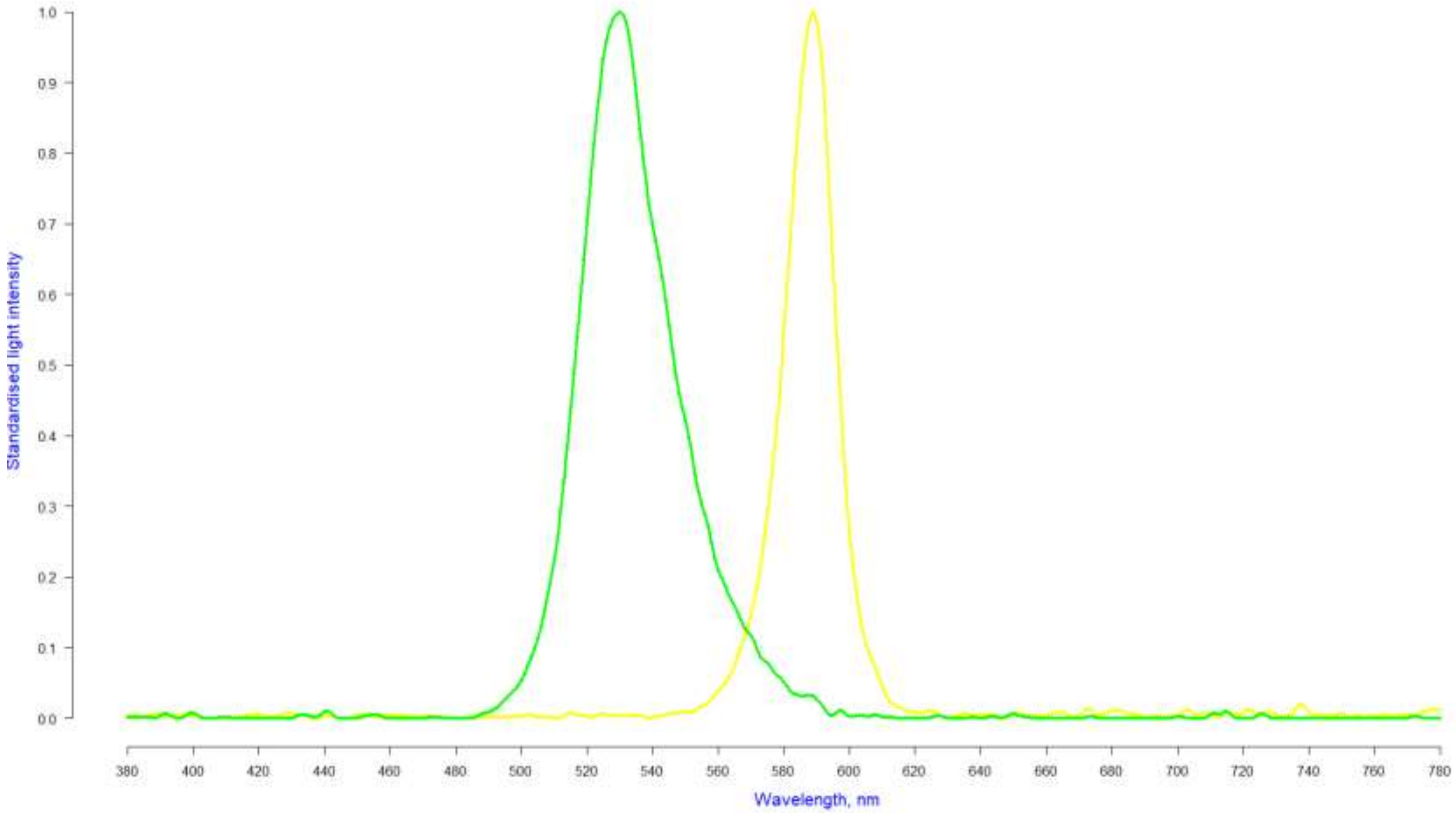

**Figure S3. Results of bootstrap analysis.**

Each diagram represents a distribution of lengths of mean vectors that were calculated using a bootstrap technique (n = 100000, see details in the main text of manuscripts, Materials and methods section): A) for birds in green light, NMF (Fig. 3A; 95 and 99 % quantiles for r mean are 0.48- 0.86 and 0.44 < r <0.94, respectively); B) for birds in green light, CMF (Fig. 3B; 95 and 99 % quantiles for r mean are 0.42 < r <0.93 and 0.38 < r <0.94, respectively. Vertical blue and red lines indicate 95 and 98 % quantiles, respectively. The red curve is a normal distribution, an orange dot is a length of the mean vector of group in each experimental condition

**A**

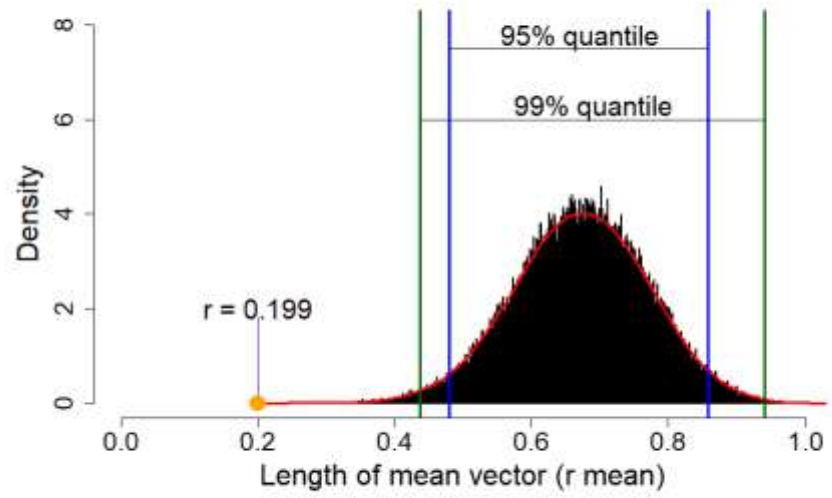

**B**

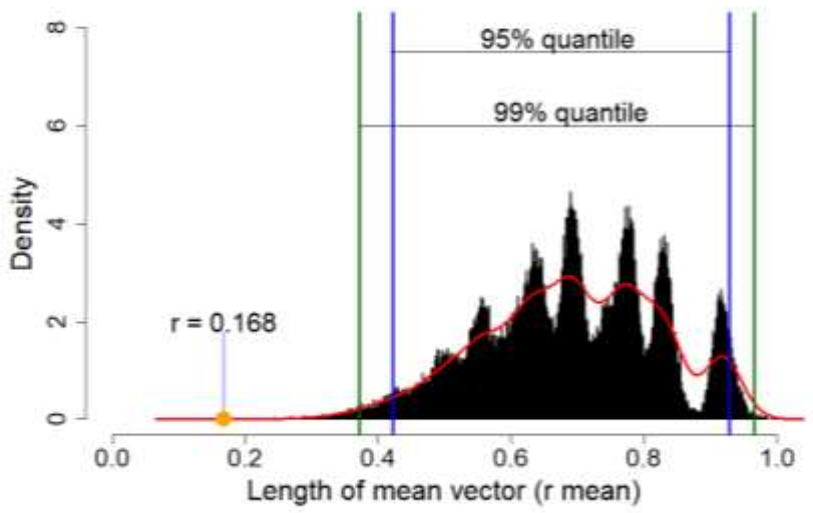
